# Antifungal effects of 3-(4-Phenyl-thiazol-2-yl)-2-thioxo-2, 3-dihydro-1H-quinazolin-4-one against *Aspergillus Species*

**DOI:** 10.1101/448704

**Authors:** Desh D. Singh, V. K. Tiwari, Rambir Singh, G L Sharam, Rajesh Dabur

**Affiliations:** Amity Institute of Biotechnology, Amity University Rajasthan, India; Department of Chemistry, Faculty of Science, Banaras Hindu University, Varanasi-221005, India; Department of Biomedical Sciences, Bundelkhand University, Jhansi-284128, India; Institute of Genomics and Integrative Biology, Mall Road, Delhi-110007; Department of Biochemistry, MDU, University Rohtak, Hariyana, India

**Keywords:** Aspergillus *fumigatus*, Antifungal activity, quinazoline, Susceptibility assay, Amphotericin B

## Abstract

Aspergillus infections have become an important health problem with the increasing number of patients. Available antifungal drugs are lack with their spectrum, toxic or immunosuppressive in nature, so that need to develop new compound with high efficacy. To evaluate antifungal efficacy of synthesized compound and to identify the protein profile of *Aspergillus fumigatus* treated with antifungal. Clinical isolates of *A. fumigatus*, *A. flavus* and *A. niger* were cultured and efficacy of compound were conducted by Disc Diffusion Assay (DDA), Microbroth Dilution Assay (MDA). Percent of spore germination inhibition assay (PSGI), Time kill analysis and toxicity assay. The culture filtrate containing secretory proteins was collected after 24 h growth and expression of downregulated proteins were identified. We developed a new and useful quinazoline derivatives expected to antifungal activity. The result of anti-Aspergillus evolution revealed that one of the 3-(4-Phenyl-thiazol-2-yl)-2-thioxo-2, 3-dihydro-1H-quinazolin-4-one (DDVKT4Q) exhibited appreciable activity. The potency of compound was found concentration of 3.125 µg/disc by disc diffusion assay (DDA) and 15.625 µg/ml. by Microbroth Dilution Assay (MDA). The compound was nontoxic up to concentration 625 µg/ml and its lysed only 35.9% of human erythrocytes, at the highest dose tested. It’s observed that the treatment of pathogen with DDVKT-4Q targeted the expression of four proteins having molecular weights 18 kDa 37 KDa and 43 KDa proteins was completely inhibited or down regulated by the compound the extra cellular. The novel compound DDVKT-4Q, having antifungal activity Can be exploited further to develop new ideal antimycotic drugs.

## Introduction

Invasive fungal infections are increasing threat to public health in last few decade, fungal infection mainly arise due to increasing immunosuppressive therapy. *Aspergillus species in*immunocompromised hosts represents a major cause of morbidity and mortality (1). The number of patients is increasing due to the transplantation for end organ disease, the most common invasive mold infection in these patients, and mortality rates in high-risk groups of myeloablative and immunosuppressive therapies for autoimmune disease, neoplastic, leukemic patient’s, hematopoietic stem cell transplant recipients and the human immunodeficiency virus/AIDS pandemic (2). *Aspergillus fumigatus* antigens are associated with asthma which amplified occurrence brings a parallel rise in the number of individuals predisposed to allergic bronchopulmonary aspergillosis, a disease associated with aberrant responses to *Aspergillus* antigens (3). Damage from *A. fumigatus*can result from fungal growth and tissue invasion or from inflammatory cells recruited to sites of infection (4). Mortality of invasive fungal infection increased up to 50% even with identification and characterization of new diagnostic techniques and novel antifungal molecules (5). The development of antifungal molecules and regular efforts are required to improve therapeutic intervention (6). In this study we report the utility of quinazoline derivative as antifungal compound. We developed a simple and convenient method for the synthesis of diverse 2-thioxo-2,3-dihydroquinazolin-4(1*H*)-ones were developed as one-pot reaction of anthranilic acid esters, primary amines, and bis (benzotriazolyl) methanethione in presence of the amidine base *DBU*(1). We evaluate the efficacy of synthesized compound and studied the protein profile of Aspergillus*fumigatus* (AF).

## Material and Methods

### Evaluation of antifungal efficacy of synthesized compounds

#### Pathogens

Clinical isolates of*A. fumigatus*, *A. flavus*and *A. niger* obtained from the Various hospitals of India was used along with the standard strains of A.*fumigatus* (ITCC 4517), *A. flavus* (ITCC 5192), *A. niger* (ITCC 5405) were used in various experiments carried out in the present study.

#### Culture of Pathogens

The pathogenic strains of *Aspergillus* were cultured for 96 h at 37^0^C. These cultures were used as the source of spores for all *in vitro* and *in vivo* experiments.

#### Preparation of Spore Suspension

Spores/conidia from the fungal colonies of 96 h cultures were adjusted to 1×10^6^spores/ml and use for further experiment.

#### Stock Solution of Standard Drug

Amphotericin B was used as the standard drug in the present study. A stock solution of standard drug was prepared in DMSO. One mg of amphotericin B was dissolved initially in 10.0 μl of DMSO and diluted with double distilled water to make up the volume to 1.0 ml. Further dilutions of the drug were made as per the need of the experimentation. Freshly prepared solutions of standard drug were used in various experiments.

### *In vitro* antifungal susceptibility assays

#### Disc diffusion assay

The disc diffusion assay (DDA) using Sabouraud dextrose agar (SDA) was performed following the method given by Yadav et al. 2005 (8).

#### Micro broth dilution assay

Susceptibility testing were subjected by following the CLSI M38-A2 broth microdilution Clinical and Laboratory Standards Institute and [CLSI]. Aspergillus species were grown overnight on Sabouraud dextrose agar (Hi-media) (9,10,11) at 35 ^o^C. Antifungal test was evaluated with RPMI-1640 medium that was buffered to pH 7.0 with 0.165 M of morpholinepropanesulphonic acid (MOPS; Sigma). Medium effects were determined by using buffered RPMI 1640 supplemented with 20.0 g of glucose per liter, Asparagine broth containing 7.0 g asparagine, 7.0 g ammonium chloride, 1.31 g potassium dihydrogen phosphate, 0.90 g sodium citrate, 10.00 g dextrose, 25.00 ml glycerol, 1.50 g magnesium sulphate, 0.30 g ferric citrate were dissolved in 1.0-liter distilled water and pH was adjusted to 7.0. Czapek Dox Broth was used Ingredients Gms / Litre containing Sucrose 30.000 Sodium nitrate 3.000 Dipotassium phosphate 1.000 Magnesium sulphate 0.500 Potassium chloride 0.500 Ferrous sulphate 0.010 Final pH (at 25°C) 7.3±0.2; Malt Extract Broth was used as Ingredients of Gms / Litre Malt extract 17.000 Mycological peptone 3.000 Final pH (at 25°C) 5.4±0.2. Mycological Broth Ingredients was in gms / Litre Papaic digest of soyabean meal 10.000 Dextrose 40.000 Final pH (at 25°C) 7.0±0.2. Fungi Kimmig broth base comprises Gms/Litre of 9.300Casein enzymic hydrolysate 4.300 Sodium chloride 11.400 Dextrose 10.000 broth, 15.000 Final pH (at 25°C) 6.5±0.2 was used and autoclaved at 15 lbs pressure (121°C) for 15 minutes. Corn Meal broth ingredients Gms/Litre Corn meal, infusion from 50.000 Agar 15.000 Final pH (at 25°C) 6.0±0.2 was used and autoclaved at 15 lbs pressure (121°C) for 15 minutes. Dilutions of antifungal agents range from 1000.0 μg/ml to 3.9032μg / ml were prepared with RPMI-1640 medium that was buffered to pH 7.0 with 0.165 M of morpholinepropanesulphonic acid (MOPS; Sigma). The drug dilutions were dispensed into round-bottomed microdilution plates (Corning Costar) and frozen at) −70C until required. The conidial inoculum suspensions were adjusted with 1×10^6^ spores in 10 μl, and diluted 1:50 in RPMI; each well was then inoculated with 100 μl, of the corresponding suspension. Various concentration of compound in the range of 1000.0 μg/ml to 3.9032μg / ml was prepared in the wells by two-fold dilution method. The wells were inoculated with 1×10^6^ spores in 10 μl of spore suspension. Appropriate control wells treated with amphotericin B or without any treatment were included in the study. The plates were incubated at 35 °C and examined visually after 48 h for the growth of *A. fumigatus* mycelia. The activity was represented as −ve if visible growth was there and +ve if medium appeared clear without any growth of mycelia of *A. fumigatus.* The lowest concentration, which inhibits the growth of*A. fumigatus*, the lowest concentration of a compound that causes a specified reduction in visible growth of an Aspergillus in broth dilution was considered as the minimum inhibitory concentration (MIC) of the compound.

#### Spore germination inhibition assay

Various concentration ranging from 1000.0 μ g/ml to 3.9032μg / ml of the DDVKT-4Q to DDVKT-4Q in 90.0 μl of culture medium were prepared in 96 well flat bottom micro-culture plates (Nune, Nunclon) by the method given by Yadav et al. 2005 (8).

### Time Kill Analysis

The assay was performed by using Dabur *et al.,* 2005 (12).

#### Toxicological evaluation of the Active compound

The *in vitro* toxicity of the compound 4 (Q) was studied by MTT assay using RAW cells.^13^

### Hemolytic assay

The assay was performed by using Yadav et al., 2005 (13).

#### To study the Protein profile of *A. fumigatus* treated with antifungal

The experiments were carried out to identify the gene/gene product (s) of A. *fumigatus* (ITCC 4517) targeted by the compound **DDVKT-4Q (7).** The pathogenic *A. fumigatus* was cultured in absence or presence of sub-lethal doses of the compound. The proteins were isolated and separated on SDS gels for comparison of protein profile to identify the gene product affected by the compound.

### Treatment of A. fumigatus in broth

A. fumigatus was grown in the presence or absence of compound in asparagine broth. A volume of 1000.0 ml of medium containing sub lethal doses of the most active compound i.e. **DDVKT-4Q** was inoculated with spores of A. fumigatus. The flasks were incubated at 37 ^°^C in BOD incubator for 24 h. The culture in medium without the compound served as appropriate controls.

#### Extraction of secretary proteins from A. fumigatus

The culture filtrate containing secretory proteins was collected after 24 h growth and transferred to a bottle containing cocktail of protease inhibitors. The filtrate was concentrated with the help of polyethylene glycol to reduce the volume by 20 times. The concentrated protein solutions were dialyzed against distilled water with change of distilled water 3 times over a period of 24 h. Both control and compound treated culture filtrates were properly labeled and protein concentration was determined.

### Protein estimation

The basic method of Smith et al (1985) described for quantification of protein by bicinchonic acid assay was used in the present study (14).

#### Sodium Dodecyl Sulphate Polyacrylamide gel Electrophoresis (SDS-PAGE)

SDS-PAGE was performed by shah et al., 2016 with slight modifications (15).

### N-Terminal Sequencing of Protein

The proteins obtained from normal culture and that treated with the purified compound were electrophoresed on 12.5% SDS polyacrylamide gels and transferred on to the polyvinylidene difluoride (PVDF) by following protocol according to Speicher et al., 2009 (16). The N-terminal amino acid sequence of the protein obtained was submitted to Blast Entrez search for sequence homology with the proteins of *A. fumigatus*and others. The results were obtained and analyzed for levels of homology.

## RESULTS AND DISCUSSION

The various test concentrations ranging from 400.0 0 μg to 1.5625 0 μg/disc of compounds were used to determine the antifungal activity by disc diffusion assay. The DDVKT-4Q disc (Fig.-1) was found to have highest activity against *A. fumigatus* at a concentration of 3.125 0 μg/ of 2-thioxo-2, 3-dihydroquinazolin-4(1H)-one derivatives.

**Fig 1.**
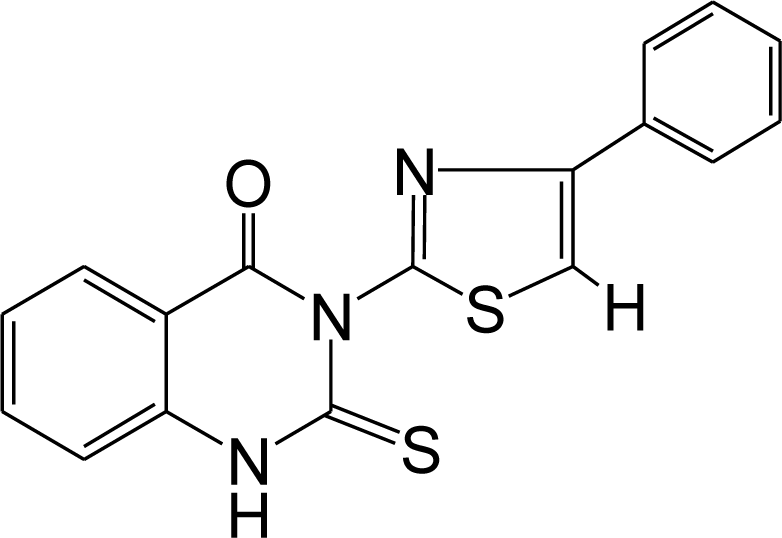
3-(4-Phenyl-thiazol-2-yl)-2-thioxo-2,3-dihydro-1H-quinazolin-4-one.

Clinical isolates of 53 strain of Aspergillus were screened to test the compound for their antifungal potential. The various test concentrations ranging from 400.0 0 μg to 1.5625 0 μg/disc of compounds were used to determine the antifungal activity by disc diffusion assay. The DDVKT-4Q was found to have highest activity against *A. fumigatus* at a concentration of 3.125 0 μg/disc (Fig.-2). Various concentration of compound in the range of 1000.0 μg/ml to 3.9032μg / ml were used MDA for synthesized compounds, MIC was found 15.625 to 31.25 µg/ml as shown in table 1 and 2 and fig 3. Diverse MIC in the group of strain tested were observed and no resistance were found against tested compound. The various test methods and culture medium can affect the efficacy of antifungal molecules. We compared the in vitro activity of compound against clinical isolates of Aspergillus species with RPMI 1640, RPMI 1640 plus 2% glucose, Yeast Nitrogen Broth (YNB) plus 0.5% glucose, Asparagine broth (AB), Czapek Dox Broth (CDB), Malt Extract (ME), Mycological Broth (MB), Sabourd Dextrose broth (SDB), antibiotic medium-3 (ABM)and Corn Meal broth (CMB) as shown in table 2. The enrichment of glucose to RPMI 1640 effect the antifungal activity does not alter the antifungal activity. The MIC antifungal molecules were observed lower in SDB, MB, and ABM 3 than RPMI then another tested medium. Amphotericin −B were observed as most active in all tested medium. The percent spore germination inhibition increased with increase in concentration of Compound used in the test. A concentration of 15.6125 µg /ml of *A. fumigatus* inhibited the growth mycelia (Fig. 4). The fungicidal activity was performed against A. fumigatus by using time kill analysis and observed that *99.8*% of loss of viability of spore after 8.5 h of incubation; however, Amp B caused 99.8 % loss of viability of spore after 4.5 h incubation, while amphotericin B caused a 99.9% loss of viability after 1 h of incubation (Fig 5). Hemolysis of the compound was compared with Amp B in % Hemolysis assay, the compound was found to be nontoxic at concentration 625 µg/ml. and its lysed only 35.9% of human erythrocytes, at the highest dose tested, that is, at the concentration of 10,000 µg/ml. Amp. B was found to be lethal to all cells at 39.05 µg/ml (Fig. 6).

**Fig 2.**
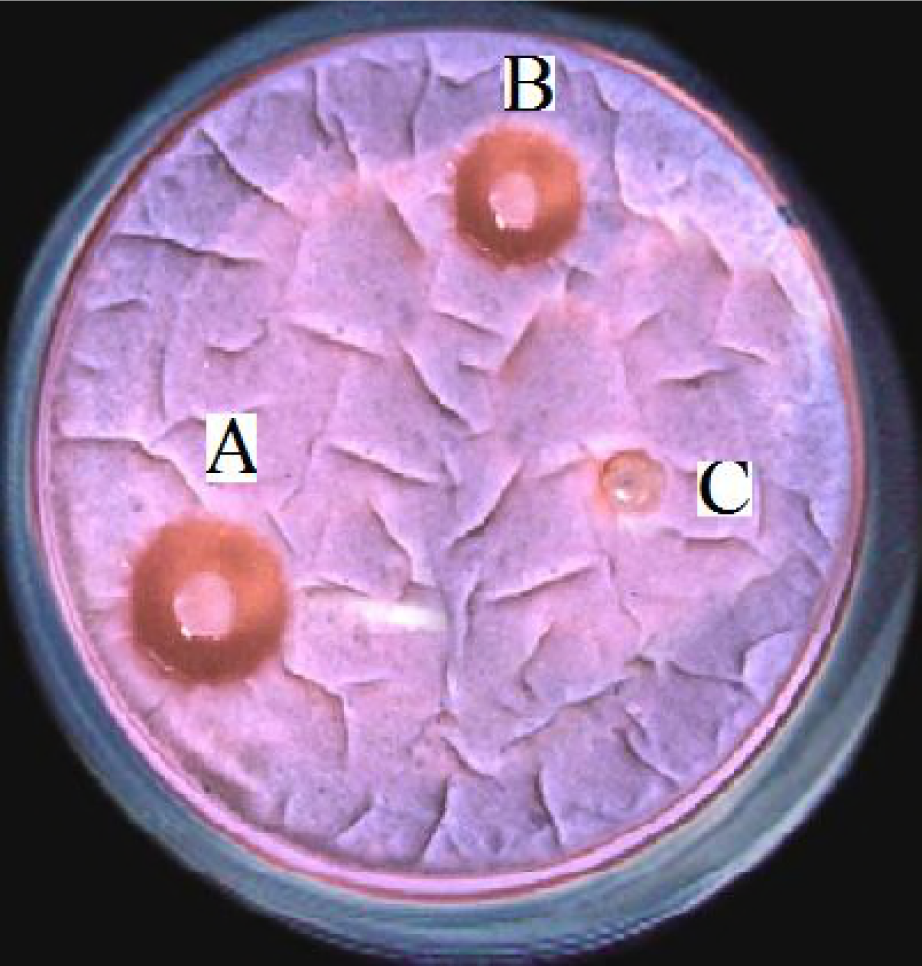
Inhibition of growth of *A. fumigatus* by DDVKT-4Q. Disc A, 3.125(μg/disc) impregnated with compound DDVKT-4Q, Disc B impregnated with Amphotericin B (1.25μg/disc) and disc C is Solvent as control.

**Fig 3.**
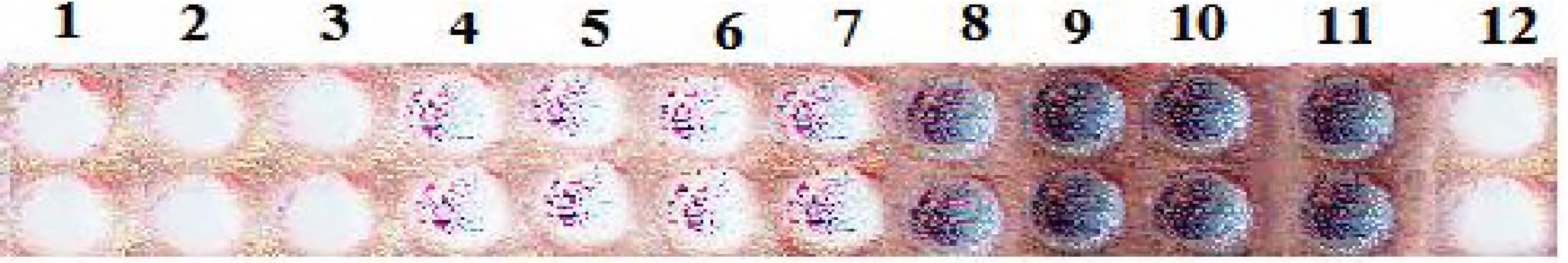
Inhibition of growth of *A. fumigatus* by DRVKT-4Q in Microbroth broth Dilution a the duplicate wells from 1-9 was inoculated with 1000 to 3.9032 μg/ml. of DDVKT-4Q. 10^th^ are normal growth and 11^th^ and 12^th^ pair of wells were negative (solvent) and positive controls (Amp. 5.0 μg/ml.). MIC was found to be 15.625 μg/ml. (lane 7^th^)

**Fig 4.**
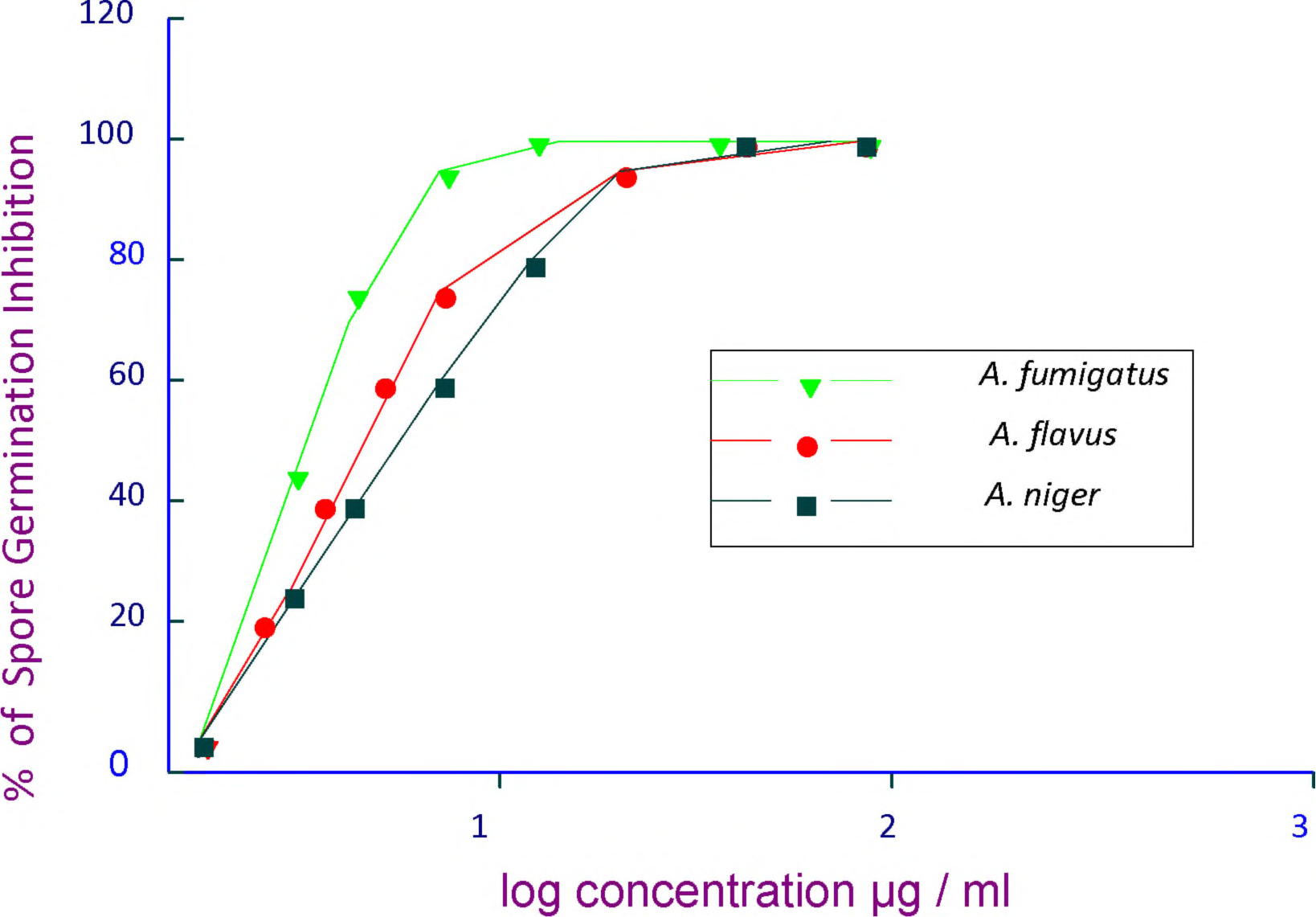
Percent spore germination inhibition by compound DDRVKT-4Q against Aspergillus species. ▾ *A. fumigatus • flavus ▪ A. niger*

**Fig 5.**
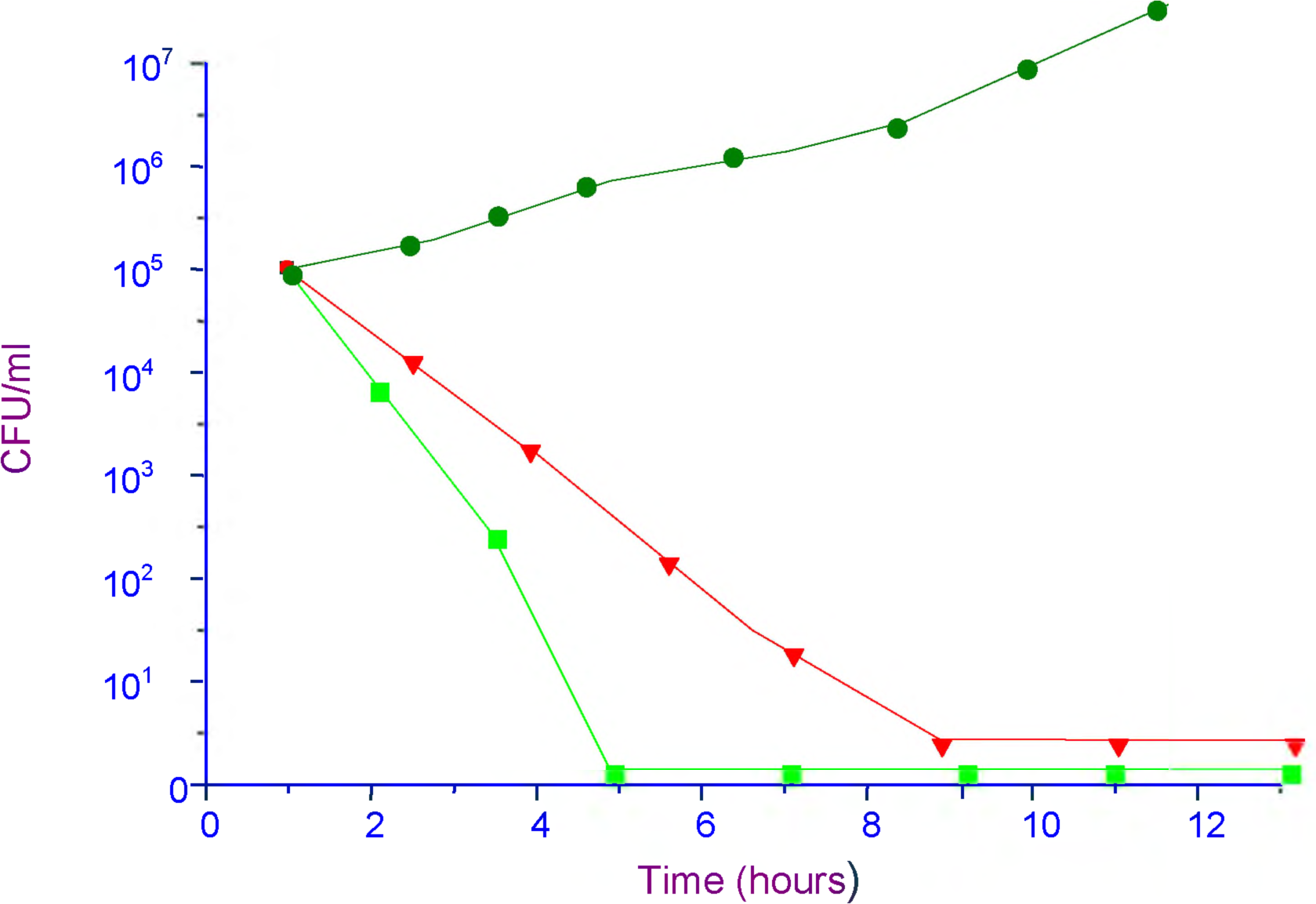
Time kills analysis of DDRVKT-4Q and amphotericin B atconcentrations equal to four times the MIC for *A. fumigatus* • drug-free control; 3.125μg/ml ▾ DDRVKT-4Q ▪ amphotericin B, 1 μg/ml.

**Fig 6.**
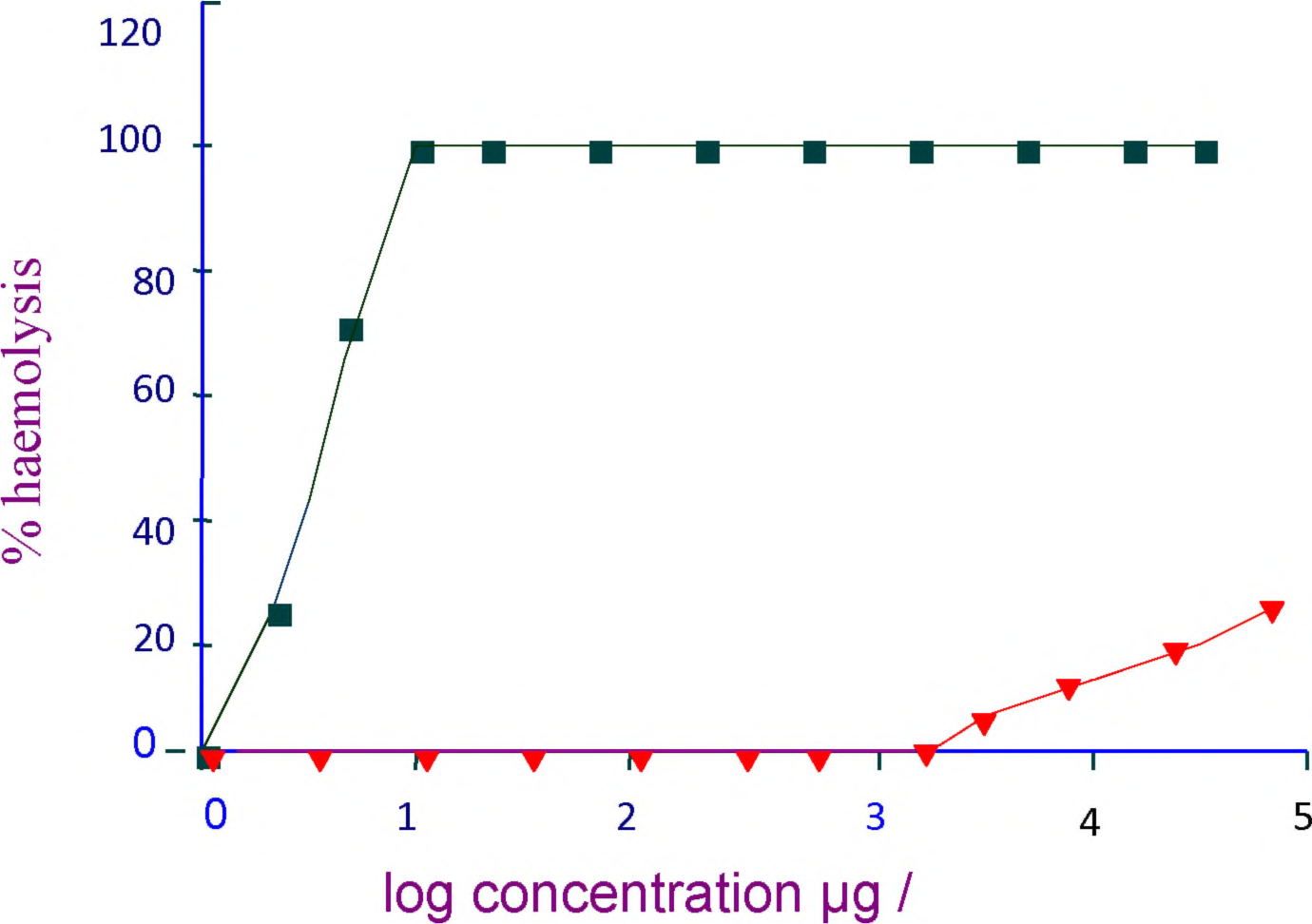
Haemolytic assay against erythrocytes using Haemolytic assay-▾ DDVKT-4Q; ▪ Amphotericin B.

**Table 1.** Comparative *in vitro* activities of amphotericin B and compound.

| Species | No. of isolates | Antifungal agent MIC (µg/ml) Compound | Antifungal agent MIC (µg/ml) Amphotericin -B |
| --- | --- | --- | --- |
| *Aspegillus fumigatus* | 20 | 15.625 | 0.5-1.0 |
| *Aspergillus flavus* | 15 | 31.25 | 0.5-1.0 |
| *Aspergillus niger* | 18 | 31.25 | 0.5-1.0 |

**Table 2.** Effect of different test media on *in vitro* activity of compound.

| Strain | No of Isolates | RPM I 1640 | RPMI 1640 plus 2% glucose | YNB plus 0.5% glucose | Asparagine broth | Czapek Dox Broth | Ma It Ext rac t Br oth | Mycologi cal Broth | Sabour d Dextro se broth | Corn Meal broth |
| --- | --- | --- | --- | --- | --- | --- | --- | --- | --- | --- |
| *A. fumigatus* | 20 | 15.625 | 15.625 | 7.815 | 15.625 | 7.815 | 15.625 | 7.815 | 7.815 | 15.625 |
| *A. niger* | 15 | 31.25 | 31.25 | 31.25 | 31.25 | 15.625 | 31.25 | 15.625 | 15.625 | 31.25 |
| *A. flavus* | 18 | 31.25 | 31.25 | 15.625 | 15.625 | 15.625 | 15.625 | 7.815 | 15.625 | 15.625 |

The asparagine broth was inoculated with spores of *A. fumigatus* in presence or absence of the test compound DDVKT-4Q. An equal amount of proteins from normal and treated culture were separated on 12.5 % polyacrylamide gel and was silver stained. The secretory proteins of untreated and DDVKT-4Q treated cultures of *A. fumigatus* after 16 h were analyzed to determine the effect of the compound on gene products of the pathogen. The protein concentration in culture filtrate was determined and equal amounts of both proteins (untreated and DDVKT-4Q treated) were run on SDS-PAGE. The treated culture was found to have less growth as compared to normal culture. This indicated that in presence of compound DDRVKT-4Q, there was less or no expression of some of the proteins during the growth of *A. fumigatus*. The expression of four proteins having molecular weights of 18 KD, 37kD, 43 KD and 70 KDa was completely inhibited or down regulated by the compound (Fig.7). These results indicated that the genes of these four proteins were the potential targets for the compound. Asp f 1 is 18 KDa major 149-amino acid protein to the ribotoxin family and secreted fungal ribonucleases whose toxicity comes from their ability to reach the cytosol via endocytosis without any receptor interaction. Once inside the host cell, ribotoxins inhibit protein biosynthesis by inactivating the ribosomes leading to cell death. The proteins of this group are cytotoxic ribonucleases that degrade a single phosphodiester bond of the 29S rRNA of eukaryotic ribosomes. We observed that the treatment of *A. fumigatus* with test compound down regulated the expression of Aspf1 ribotoxins. Reduced expression of this protein will decrease the pathogenicity of the fungus. The mechanism for down regulation of this protein need to be explored further.

**Fig 7.**
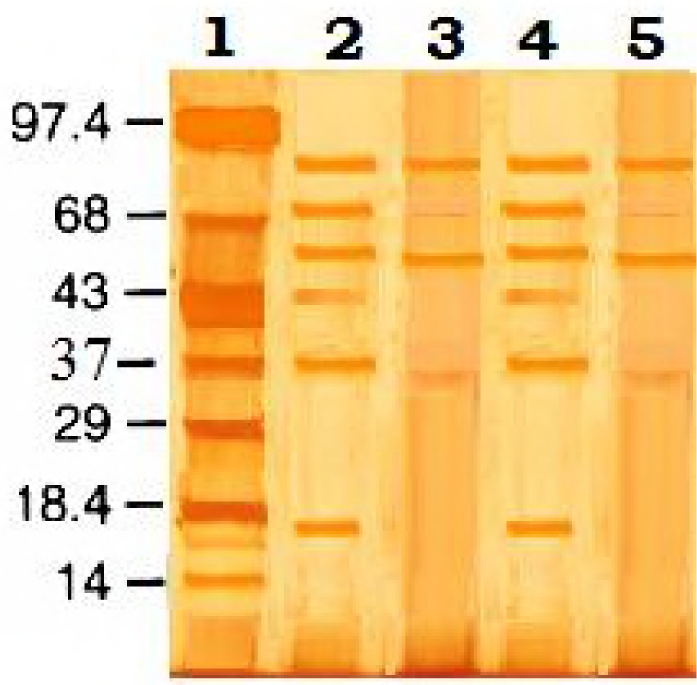
Profile of secretory proteins (12.5% acrylamide SDS gel) of normal and compound treated cultures of A. fumigatus. Lane 1: Marker lane, 2 and 4 Proteins from untreated cultures. Lane 3 and 5: Proteins from DDVKT −4Q treated cultures.

The second down regulated protein was found to be 37-kDa IgE binding epitopes of *A fumigatus. S*everal earlier studies have reported that Aspf 2 having cysteine residues with sulfhydryl groups in their side chains are involved in interchain or intrachain disulfide bond formation with IgE Ab. Asp f 2 residual interactions occur due to IgE Abs with epitopes involving amino acids cysteines. This antigen is responsible for induce Type I and Type III hypersensitivity reactions leading to allergic bronchopulmonary aspergillosis (ABPA) in immunocompetent host. The compound DDRVKT-4Q could be a potential role in the A*. fumigatus*-associated allergic diseases.

Third protein was found to be a 43 kDa metalloprotease have been reported to be involved in the degradation of lung structural material, metalloprotease was able to hydrolyze elastin, and its elastinolytic activity. There is an 18 amino acid long signal peptide metalloprotease protease of *A. fumigatus*may be the possible target of compound.

Galactomannan (GM) is 70-kDa GMP2 secreted during growth in culture, cell wall-linked GMs have a similar chemical organization composed of a linear core with a repeating tetra mannose unit (2M-6M-2M-2M) with side chains of β-1,5- glucofuranose residues with an average degree of polymerization of 4 attached to the β-1,5- glucofuranose −1,2-linked mannose residues of the mannan chain. Therefore, it seems that GM protein is involved in membrane integrity which are completely down regulated when treated with compound, could be major target for drug development against Aspergillosis.

The compound 3-(4-Phenyl-thiazol-2-yl)-2-thioxo-2, 3-dihydro-1H-quinazolin-4-one, having antifungal activity can be exploited further to develop new ideal antimycotic drugs.

## Conclusion

New antifungal agents are needed due to the importance of fungal infections in compromised patients, limitations of currently available antifungal agents regarding their spectra of activity and toxicities, and the increasing prevalence of pathogens resistant to the current antifungal agents. Tested compound has shown maximum fungicidal activity against the most clinically relevant *A. fumigatus, A niger and A. flavus*. The results clearly indicate that the compound 3-(4-Phenyl-thiazol-2-yl)-2-thioxo-2,3-dihydro-1H-quinazolin-4-one is potential lead compound for Aspergillosis therapy. This compound also has low cytotoxicity; therefore, it may not only be a therapeutically useful antifungal agent but also a model compound to develop good fungicidal compounds.

## ACKNOWLEDGEMENTS

The author gratefully acknowledges Institute of Genomics and integrative Biology, Delhi University camus, Mall road delhi, Dr. A K. Mohanty, Proteomics and Structural Genomics lab, Animal Biotechnology Division, NDRI, Karnal India; Prof. Amita Jain, Department of Microbiology King George’s Medical University, Lucknow, India, Institute of Biomedical sciences, Bundelkhand University, Jhansi, India; Department of Chemistry, Banaras Hindu University, Varanasi, India.

